# MitoStores: Chaperone-controlled protein granules store mitochondrial precursors in the cytosol

**DOI:** 10.1101/2022.08.02.502474

**Authors:** Lena Krämer, Niko Dalheimer, Markus Räschle, Zuzana Storchová, Jan Pielage, Felix Boos, Johannes M. Herrmann

## Abstract

Hundreds of mitochondrial precursor proteins are synthesized in the cytosol and imported into mitochondria in a post-translational reaction. The early processes associated with mitochondrial protein targeting remain poorly understood. Here we show that in baker’s yeast, the cytosol has the capacity to transiently store matrix-destined precursors in dedicated deposits which we named MitoStores. MitoStores are strongly enhanced when protein import into mitochondria is competitively inhibited by a clogging of mitochondrial import sites, but also formed under physiological conditions when cells grow on non-fermentable carbon sources. MitoStores are enriched for a specific subset of nuclear encoded mitochondrial proteins, in particular those containing N-terminal mitochondrial targeting sequences. MitoStore formation is controlled by the heat shock proteins Hsp42 and Hsp104, potentially to suppress the toxic potential of accumulating mitochondrial precursor proteins. Thus, the cytosolic protein quality control system plays an active role during early stages in mitochondrial protein targeting by the coordinated and localized sequestration of mitochondrial precursor proteins.

**Summary:** The yeast cytosol can deposit precursors of mitochondrial proteins in specific granules called MitoStores. MitoStores are controlled by the cytosolic chaperone system, in particular by Hsp42 and Hsp104. MitoStore formation suppresses the toxicity arising from non-imported mitochondrial precursor proteins.

## Introduction

Mitochondria consist of hundreds of different proteins that are synthesized in the cytosol and subsequently imported into mitochondria by translocases in the outer and inner membrane (Chacinska et al., 2009). In nerve, muscle or brown adipose cells, but also in respiring yeast cells, mitochondrial precursors represent a considerable fraction of all nascent protein chains. Only a minor fraction of the nuclear encoded mitochondrial proteins is synthesized on the mitochondrial surface (Williams et al., 2014). Owing to the post-translational import mode of most mitochondrial proteins (Wienhues et al., 1991), cytosolic precursors explore the cytosol, strain on the cytosolic chaperone system (Deshaies et al., 1988; Jores et al., 2018) and pose a threat to cellular proteostasis (Nowicka et al., 2021; Schäfer et al., 2021; Sorrentino et al., 2017; Wang and Chen, 2015; Weidberg and Amon, 2018; Wrobel et al., 2015). Mitochondrial proteins of the outer membrane, the intermembrane space, the inner membrane and the matrix differ in their import routes and targeting signals; in the following we refer to the cytosolic native forms of all of these proteins as precursor proteins or precursors (Chacinska et al., 2009).

Eukaryotic cells carefully sense the levels of mitochondrial precursor proteins in the cytosol and respond to accumulating mitoprotein levels with adaptations in gene expression which are summarized with the umbrella term mitoprotein-induced stress response (Boos et al., 2020). In animal cells (in particular those of the nematode *Caenorhabditis elegans*) this is achieved via dedicated transcription factors containing a mitochondrial targeting sequence; under conditions which prevent an unconstrained uptake of these proteins into mitochondria, they enter the nucleus and launch specific transcriptional responses. This <u>U</u>nfolded <u>P</u>rotein <u>R</u>esponse of <u>m</u>itochondria (UPR^mt^) increases the import and folding capacity of mitochondria and results in the expansion of the mitochondrial network in general (Aras et al., 2020; Fiorese et al., 2016; Labbadia et al., 2017; Nargund et al., 2012; Shpilka et al., 2021; Xin et al., 2022).

In yeast, no specific transcription factors were identified, but the general accumulation of non-imported precursor proteins launches the <u>U</u>nfolded <u>P</u>rotein <u>R</u>esponse <u>a</u>ctivated by <u>m</u>istargeting of proteins (UPRam), which increases the capacity of the cytosolic ubiquitin-proteasome system (UPS) by an upregulation of constituents of the degradation machinery and by stimulating their assembly (Boos et al., 2019; Wrobel et al., 2015). This upregulation of the proteasome is mediated by the transcription factor Rpn4 (Boos et al., 2019). Rpn4 serves as the master regulator of proteasome levels in baker’s yeast and is tightly controlled by the proteasome; if the proteasome system is overstrained, Rpn4 accumulates and induces a 2-4 times upregulation of components of the ubiquitin-proteasome system compared to non-stressed conditions (Metzger and Michaelis, 2009; Xie and Varshavsky, 2001). Therebys, non-imported mitochondrial precursor proteins are under tight surveillance of the proteasome in order to prevent an overload of the mitochondrial protein import system (Fig. 1A).

**Fig. 1.**
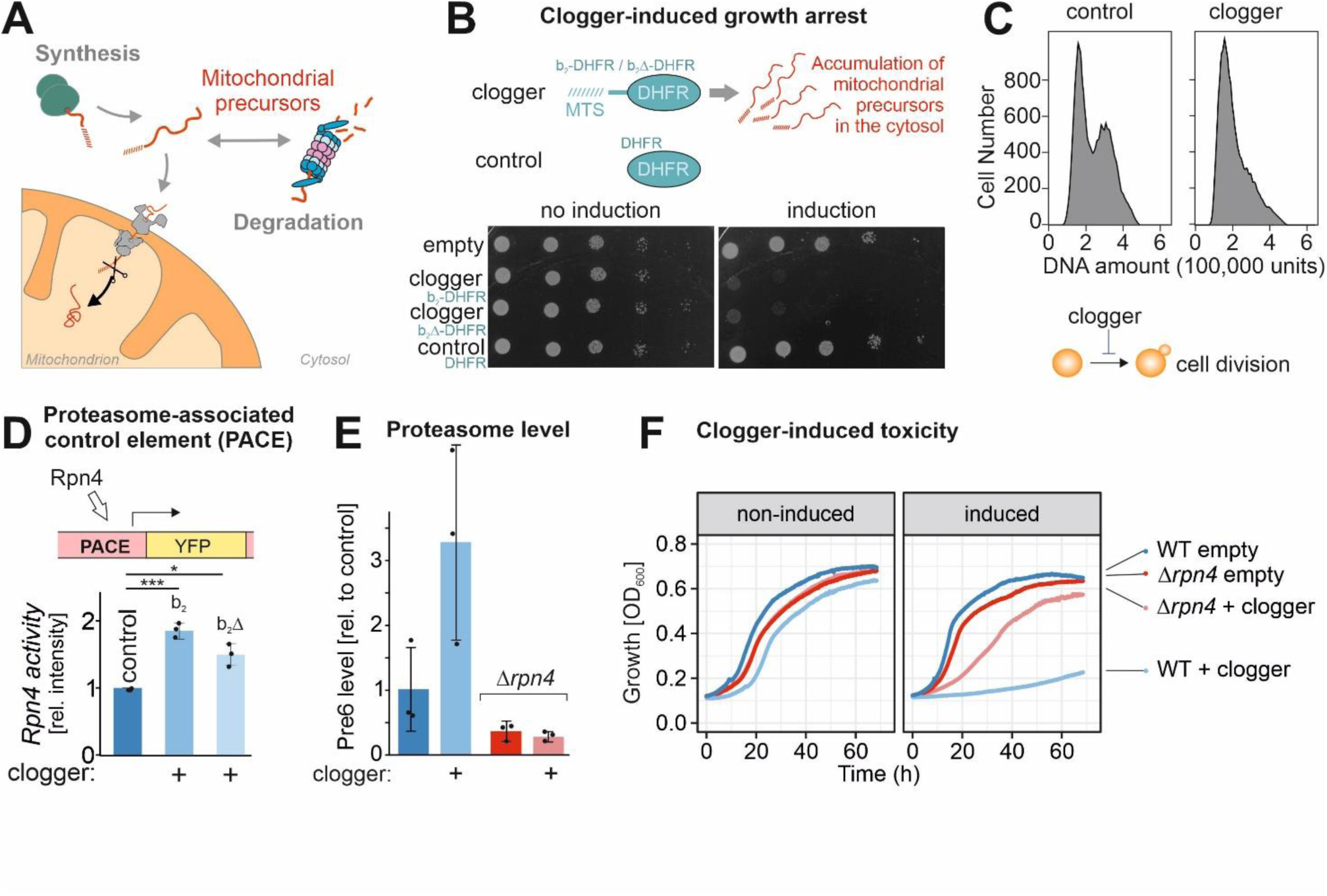
Rpn4-mediated proteasome induction represents one arm of the mitoprotein-induced stress response. (**A**) Mitochondrial protein import is under surveillance of the proteasome. (**B**) Competitive inhibition of mitochondrial protein import by clogger proteins inhibits cell growths. Yeast cells expressing clogger proteins (cytochrome *b*_2_-DHFR and cytochrome *b*_2_*Δ*_19_-DHFR) or cytosolic DHFR for comparison under control of the galactose-inducible promoter were grown to mid-log phase on lactate medium. Ten-fold serial dilutions were dropped on lactate (no induction) or lactate with 0.5% galactose medium (induction). MTS, matrix-targeting signal. (**C**) After expression of the clogger *b*_*2*_-DHFR or the DHFR control, cellular DNA was stained with propidium iodide and DNA content was assessed by flow cytometry. Please note that the first peak shows cells in G1 phase, and the second peak represents cells in the G2 phase. (**D**) Rpn4-driven gene induction was measured using a reporter that expressed the yellow fluorescence protein under control of a PACE element (Boos et al., 2019). (**E**) Levels of the proteasome protein Pre6 were detected upon expression of cytosolic DHFR or clogger for 4.5 h by Western blotting and quantified from three replicates. (**F**) Cells of the indicated strains were grown to log phase and diluted in lactate (no induction) or lactate with 0.5% galactose (induction) to 0.1 OD. Growth was monitored upon constant agitation at 30°C.

Cellular protein homeostasis, called proteostasis, is of central relevance for cellular functionality and organismal health (Labbadia and Morimoto, 2015). Proteostasis relies on the dynamic interplay of two closely cooperating systems, the network of molecular chaperones facilitating protein folding (Hipp et al., 2019) and the cellular protein degradation systems, particularly the ubiquitin-proteasome system (UPS) (Dikic, 2017; Varshavsky, 2017). Failure of either of these systems results in the accumulation of misfolded proteins, a characteristic hallmark of aging and of numerous diseases, including but not restricted to neurodegenerative amyloid disorders such as Alzheimer’s or Huntington’s disease (Hartl, 2017; Pilla et al., 2017).

When chaperones and the proteasome are overstrained, cells use a second line of defense: to suppress their cellular toxicity, misfolded proteins can be spatially sequestrated at dedicated locations. Several types of quality control compartments were previously defined based on their intracellular location and protein composition (Sontag et al., 2017). In the baker’s yeast *Saccharomyces cerevisiae*, sudden shifts to high temperature, proteasome impairments, or defects in chaperone function lead to a transient accumulation of Q-bodies (also referred to as CytoQs or stress granules) which, upon prolonged stress conditions, can coalesce into larger JUNQs and INQs (juxtanuclear and intranuclear quality control compartments, respectively) on both sides of the nuclear envelope (Escusa-Toret et al., 2013). Whereas the proteins of Q-bodies, JUNQs and INQs are rapidly released and degraded once stress conditions are relieved, insoluble protein deposits (IPODs), for example those formed upon expression of polyQ-expanded huntingtin, are highly stable and pose a severe threat to cellular function (Gruber et al., 2018; Mogk et al., 2018). In addition, different, less defined types of protein aggregates were observed on the surface of mitochondria and of the endoplasmic reticulum (ER)(Andreasson et al., 2019; Zhou et al., 2014). Recent studies suggest that non-imported mitochondrial precursor proteins contribute to the formation of cytosolic aggregates (Nowicka et al., 2021; Schlagowski et al., 2021; Xiao et al., 2021).

In this study, we elucidated the consequences of competitive inhibition of mitochondrial protein import in an Rpn4-deficient mutant in which the proteasome cannot be upregulated. To our surprise, we observed that these mutants escape the mitoprotein-induced growth arrest and tolerate the accumulation of mitochondrial precursors even better than wild type cells. Apparently, these strains employ an alternative mechanism to cope with non-imported protein precursors: they form cytosolic granules that we named MitoStores. We show that these structures contain a specific subset of mitochondrial precursor proteins and store them transiently until the stress is resolved. Thus, in contrast to previous assumptions (Chen and Douglas, 1987; Neupert and Herrmann, 2007), the cytosol has the capacity to store mitochondrial precursor proteins in dedicated storage granules that are controlled by the cytosolic chaperone system.

## Results

### Cells lacking Rpn4 can bypass the clogger-induced growth arrest

Overexpression of slowly imported precursor proteins, so-called cloggers, causes growth arrest of yeast cells by activating a block of cell division (Fig. 1B, C). The accumulation of mitochondrial precursor proteins in the yeast cytosol (Fig. S1A) induces an Rpn4-mediated induction of genes that are under control of a proteasome-associated control element, PACE (Boos et al., 2019; Zöller et al., 2020). Accordingly, expression of clogger proteins destined to the mitochondrial matrix or the IMS (b_2_Δ-DHFR or b_2_-DHFR, respectively; here driven by a galactose-inducible promoter) increase the level of the proteasome system in an Rpn4-dependent fashion (Fig. 1D, E, S1B, C).

However, to our surprise we observed that Rpn4 deletion mutants showed a markedly increased clogger resistance. Clogger expression in *Δrpn4* cells was better tolerated and these cells grew efficiently when the mitochondrial import system was challenged by clogger expression or by inhibition of the respiratory chain (Fig. 1F, S1D). In contrast, deletion of Rpn4 did not increase the resistance to the expression of aggregation-prone model proteins in the cytosol such as the huntingtin-derived poly glutamine protein Q97-GFP, luciferase^R188Q,R261Q^ or Ubc9^Y68L^ (Fig. S1E, S3A,B). Hence, ablation of the master regulator of the ubiquitin-proteasome system renders cells resistant against mitochondrial stress but not against proteotoxic stress in general.

### *Δrpn4* cells respond to clogger induction by upregulation of Hsp42 and Hsp104

How do *Δrpn4* cells escape the mitoprotein-induced toxicity? First, we tested whether Rpn4 influences the expression of the clogger protein. To this end, we expressed the clogger protein for 4.5 h and then switched cells back from galactose to lactate to monitor clogger stability. As expected, the reduced proteasome capacity in *Δrpn4* cells impairs the proteolytic removal of the clogger protein from mitochondria (Fig. 2A) and thereby even further increases the accumulation of mitochondrial precursor proteins in the cytosol (Fig. S1A). Thus, yeast cells normally respond to mitoprotein-induced stress by the upregulation of the proteasome system; however, if this response is prevented, there apparently exists a second line of defense that even more efficiently alleviates the toxicity arising from mitochondrial precursor proteins in the cytosol.

**Fig. 2.**
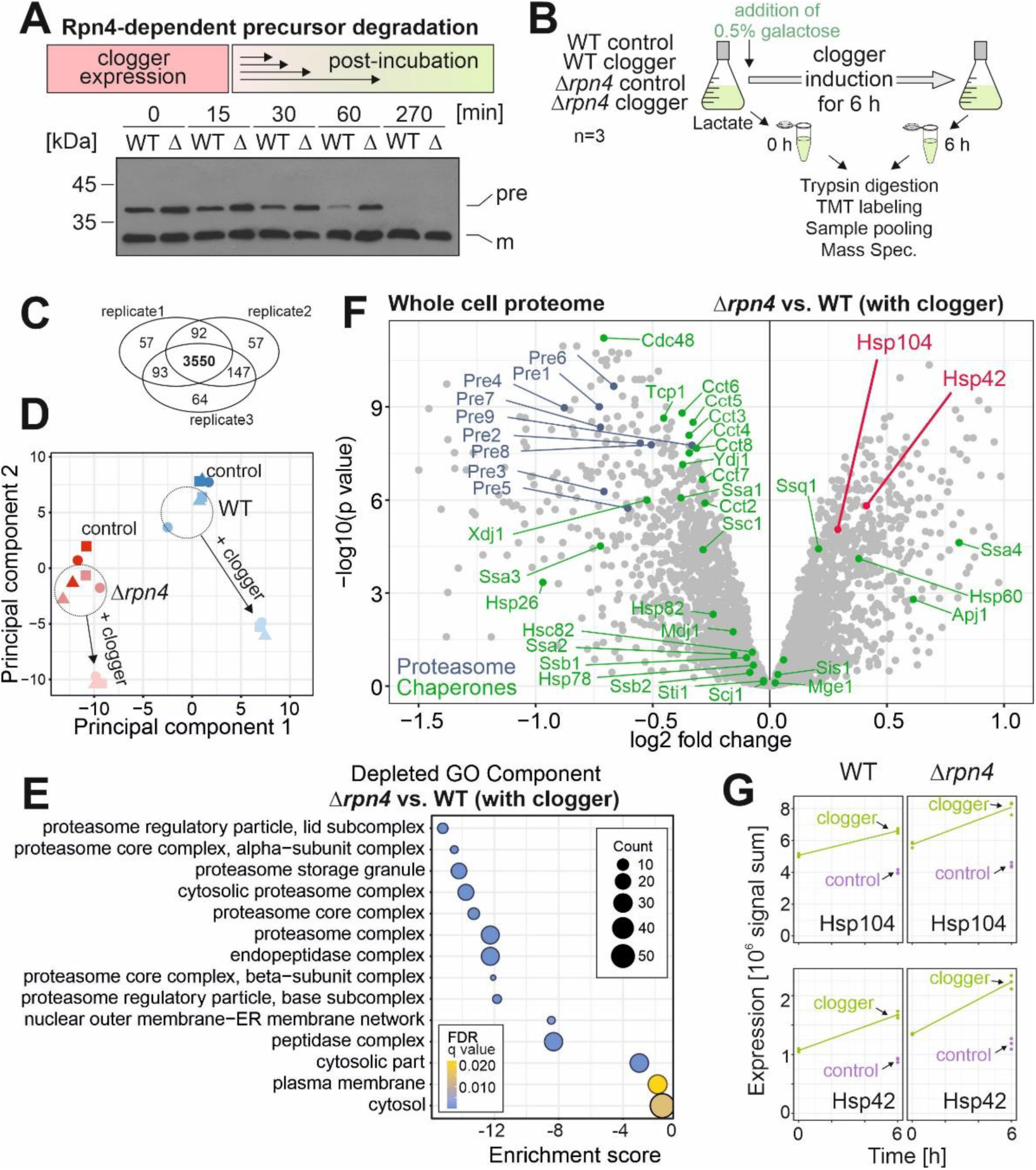
Competitive inhibition of mitochondrial protein import in *Δrpn4* cells remodels the cytosolic chaperone system. (**A**) The clogger was expressed in wild type (WT) and *Δrpn4* (*Δ*) cells for 4.5 h. Medium was exchanged for a non-inducing lactate medium. Precursor (pre) and mature (m) forms of the b_2_*Δ*-DHFR clogger were visualized by Western blotting using a DHFR-specific antibody. (**B**) The clogger was induced for 6 h before cellular proteomes were measured by mass spectrometry. **(C)** 3,550 proteins were measured in all samples of the three replicates. (**D**) Principal component analysis. Presence of Rpn4 and clogger expression caused specific changes in the proteome. The different shapes of the datapoints indicate the three different biological replicates. (**E**) GO Term Analysis was done using the GOrilla tool (http://cbl-gorilla.cs.technion.ac.il/). The absence of Rpn4 caused a broad reduction of the components of the proteasome-ubiquitin system. The number of proteins in the indicated GO Components are stated as “Counts”. (**F**) Comparison of the proteome of wild type and *Δrpn4* cells after clogger expression for 6 h. Positions of proteins of the proteasome and chaperone systems are indicated in blue and green, respectively. Please note that Hsp104 and Hsp42 show a higher abundance in *Δrpn4* cells. See Table S1 for details. (**G**) Signals of Hsp104 and Hsp42 before and after clogger induction in wild type and *Δrpn4* cells.

To identify the underlying molecular mechanism, we compared the cellular proteomes of wild type and *Δrpn4* cells before and upon clogger induction (Fig. 2B) using isobaric mass tag TMT) labeling and quantitative mass spectrometry (Dayon et al., 2008). More than 3,800 proteins were quantified in at least two out of three replicates across all samples (Fig. 2C, supplemental Table S1). This showed that the proteomes of wild type and *Δrpn4* cells considerably differ already before clogger expression (Fig. 2D) and that the loss of Rpn4 leads not only to reduced levels of proteasome proteins but also to lower amounts of proteins of the mitochondrial respiratory chain (Fig. 2E, S2A). Our observations confirmed that Rpn4 is crucial for the induction of the proteasome system and clogger induction did not bypass the Rpn4 relevance for proteasome upregulation (Fig. S2B), again suggesting that *Δrpn4* cells employ a proteasome-independent mechanism to deal with cytosolic precursor proteins.

We noticed that while the levels of most chaperones were unaffected or even reduced in *Δrpn4* cells, the disaggregase Hsp104 and the small heat shock protein Hsp42 were considerably upregulated (Fig. 2F, G), potentially as a proteasome-independent protection against accumulating mitochondrial precursor proteins.

### Hsp104 binds to cytosolic granules containing mitochondrial precursor proteins

Hsp104 is a hexameric chaperone in the yeast cytosol, which binds to misfolded or aggregated proteins and disentangles them in an ATP-dependent manner (Gates et al., 2017; Sanchez and Lindquist, 1990). Thereby, it binds different types of cellular aggregates (Sontag et al., 2017), including transiently forming benign condensates of dynamic nature (Yoo et al., 2022) but also insoluble protein deposits (IPODs) that pose a threat to cellular functionality (Gruber et al., 2018; Mogk et al., 2018). Consistent with previous studies (Kaganovich et al., 2008; Ruan et al., 2017), we found that fusion proteins of Hsp104 or an Hsp104^Y662A^ trapping mutant with GFP are well tolerated by yeast cells and serve as powerful tools to visualize condensates formed by the aggregation-prone model proteins luciferase^R188Q,R261Q^ and Ubc9^Y68L^ (Fig. S3A-F)

In glucose-grown wild type cells, Hsp104-GFP was evenly distributed throughout the cells whereas upon growth in non-fermenting lactate-based media cells showed a small number of aggregates (4 on average) (Fig. 3A). Upon clogger induction, the number of aggregates per cell did not increase (Fig. S4A), but the mean diameters of Hsp104-GFP-bound foci increased significantly 1.7-fold (545 to 915 nm) (Fig. 3A). A comparable increase (382 to 724 nm) was also observed in *Δrpn4* cells, which generally contained more aggregates (7.5 on average).

**Fig. 3.**
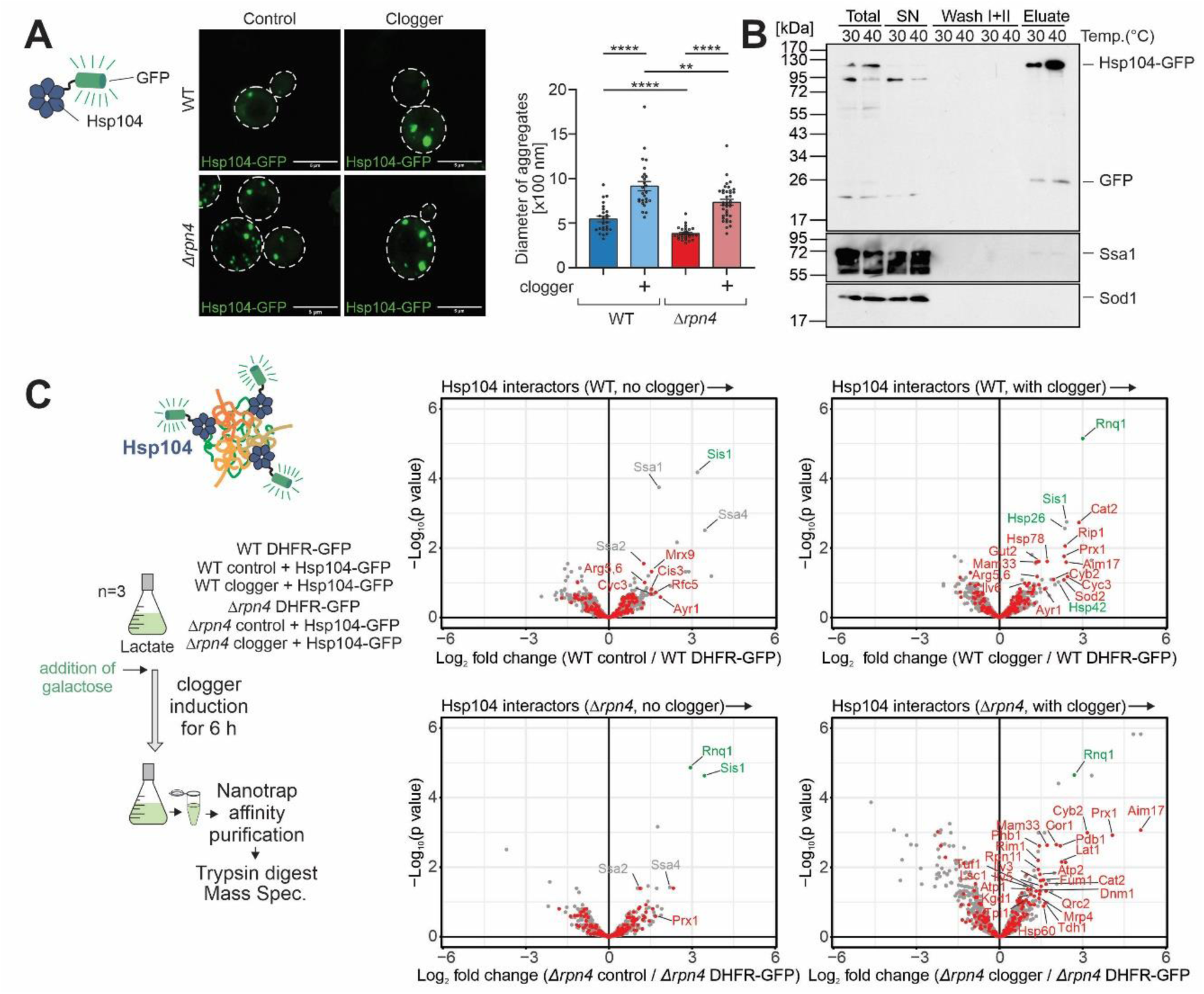
Mitoprotein-induced stress induces the formation of MitoStores in the cytosol. (**A**) Cytosolic DHFR (control) or the clogger protein *b*_*2*_-DHFR were expressed in wild type and *Δrpn4* cells for 4.5 h. The distribution of constitutively expressed Hsp104-GFP was visualized by fluorescence microscopy. Diameter of Hsp104-GFP-bound aggregates were quantified. ** p ≤ 0.01, **** p ≤ 0.0001. See also Fig. S4A for quantification of the number of aggregates per cell. (**B**) Hsp104-GFP was purified on nanotrap sepharose in wild type cells that were grown at the indicated temperatures. SN, supernatant representing the non-bound fraction. The signals for Ssa1 and Sod1 are shown for control. **(C)** Hsp104-GFP or DHFR-GFP were co-expressed with clogger or the DHFR control in wild type and *Δrpn4* cells. Cells were lysed and the extracts were subjected to nanotrap sepharose chromatography. Purified proteins were analyzed by tandem mass spectrometry. Mitochondrial proteins are highlighted in red, specific chaperones and Rnq1 are indicated in gray or green. The Hsp104 data point was omitted for better scaling of the figure. See Table S2 for details.

To identify the content of the Hsp104-GFP-bound structures, we developed a purification procedure by affinity chromatography with nanotraps after gentle lysis of yeast cells (Fig. 3B). For comparison of the Hsp104-GFP pulldown, we used isolations of a cytosolic DHFR-GFP fusion protein since this soluble and well-behaved protein accumulated at similar levels as Hsp104-GFP in the yeast cytosol (Fig. S4B) and identified the co-eluted proteins by quantitative proteomics (Fig. 3C, S4C, D).

Established interactors of the Hsp104 disaggregase such as the yeast prion protein Rnq1 or the J-domain protein Sis1 (Higurashi et al., 2008; Ho et al., 2019; Krobitsch and Lindquist, 2000; Kryndushkin et al., 2011; Wyszkowski et al., 2021) were pulled down with the Hsp104-GFP under all conditions tested. Upon clogger induction, a large number of mitochondrial proteins were co-isolated with Hsp104, indicating that the increase in size of the Hsp104-bound granules is at least in part due to the association of mitochondrial proteins with these structures. In the absence of Rpn4, this association was even further increased indicating that precursor proteins that escape proteolytic degradation in the cytosol can be incorporated into dedicated structures. These structures are reminiscent to structures that were previously observed and sometimes described as a specific type of stress granules, Q bodies or cytoQs (Böckler et al., 2017; Grousl et al., 2018; Miller et al., 2015; Nowicka et al., 2021; Shakya et al., 2021; Xiao et al., 2021). Owing to the striking acquisition of nuclear encoded mitochondrial proteins in these structures, we termed them MitoStores.

### MitoStores contain proteins with mitochondrial presequences

Next, we analyzed the Hsp104-associated proteins in more detail (Fig. 4A). To this end, we listed proteins that were considerably enriched in the Hsp104-GFP pull-downs (in comparison to the DHFR-GFP control). Many mitochondrial proteins were found as clients of Hsp104 both in the absence or presence of Rpn4. Examples of this group are Aim17, Mam33, Prx1, Ilv3, Arg5,6, Sod2 or Rip1. These proteins apparently populate MitoStores irrespective of the proteasomal activity. However, another large group of mitochondrial proteins were only found in MitoStores when the upregulation of proteasomal activity was prevented by deletion of Rpn4. These proteins, including Pdb1, Atp14, Tdh2, Cox8, Cox13, or Mrp3, are presumably tightly controlled by the proteasome and only found as clients of Hsp104 in *Δrpn4* cells. Interestingly, several of these proteins were found recently in a proteome-wide screen for mitochondrial proteins that accumulate in the cytosol when mitochondrial import is blocked by treatment with an uncoupler of the membrane potential (Shakya et al., 2021).

**Fig. 4.**
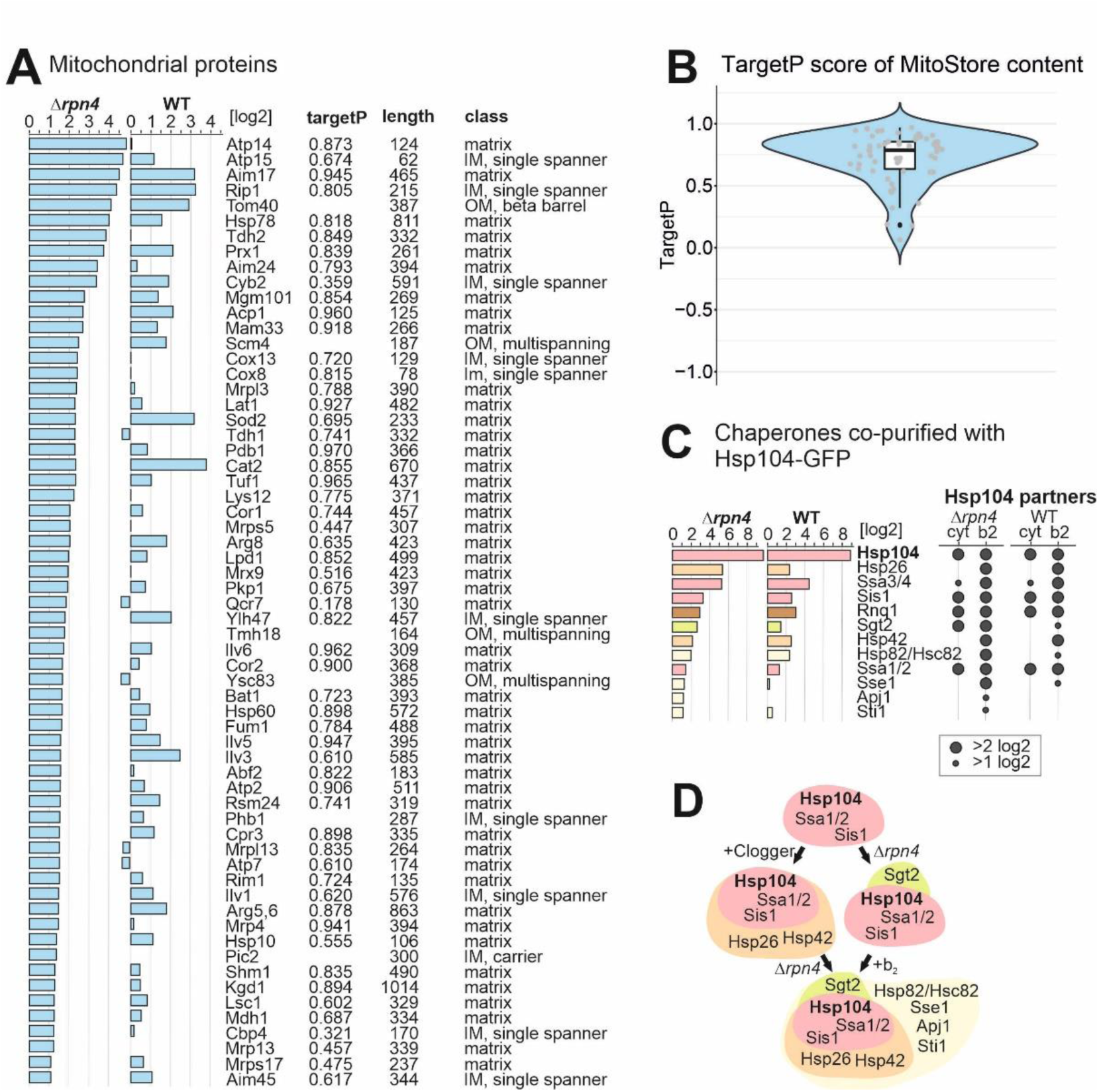
MitoStores contain mitochondrial proteins with N-terminal matrix targeting sequences. (**A**) Mitochondrial precursor proteins co-purified with Hsp104-GFP upon clogger expression in *Δrpn4* and wild type cells. Proteins were listed according to their enrichment in the Hsp104-GFP sample in comparison to the DHFR-GFP control. Scores for matrix targeting probabilities (targetP) (Emanuelsson et al., 2000) and mitochondrial location of the proteins are shown. IM, inner membrane; OM, outer membrane. TargetP values were calculated for matrix and presequence-containing IM proteins. (**B**) MitoStore clients are characterized by high probability scores for mitochondrial targeting. (**C, D**) Chaperones co-isolated on the nanotrap beads with the Hsp104-GFP protein (in comparison to DHFR-GFP) in the presence of the clogger or the control. Hsp26, Hsp42 and Sse1 are coisolated with Hsp104 in the presence of clogger, but not upon expression of the DHFR control.

We noticed that almost all mitochondrial proteins found in the MitoStores contained matrix-targeting sequences resulting in high scores in mitochondrial targeting prediction programs such as targetP (Emanuelsson et al., 2000)(Fig. 4A, B). This is surprising as only about 60% of all mitochondrial proteins contain N-terminal matrix-targeting sequences which are absent from many proteins of the outer membrane, the IMS and the inner membrane (Morgenstern et al., 2017; Vögtle et al., 2009). Apparently, the presence of mitochondrial targeting sequences supports the incorporation of clients into MitoStores, either by a specific uptake mechanism or by the kinetic properties of their cytosolic targeting (Fig. 4B). Thus, Hsp104 particularly binds to proteins with mitochondrial presequences, whereas proteins of the outer membrane, the IMS or carrier proteins of the inner membrane were considerably underrepresented. Also, several proteins of the endomembrane system were recovered with Hsp104, however, by far not as many as mitochondrial proteins (Fig. S4E).

Several factors of the cytosolic chaperone system were coeluted with Hsp104-GFP and some of these proteins were strongly enriched upon clogger induction (Fig. 4C, D, S4F). Most strikingly, the small heat shock proteins Hsp26 and Hsp42 coeluted with Hsp104-GFP once mitochondrial precursors accumulated in the cytosol, suggesting that these small heat shock proteins (Haslbeck et al., 2020; Mogk and Bukau, 2017) are MitoStore constituents. Other proteins, such as the prion protein Rnq1 were recovered with Hsp104-GFP independently of clogger expression and Rnq1 therefore might be part of another Hsp104-bound complex. Rnq1 was also not necessary for MitoStore formation (Fig. S5B).

### Mitochondrial precursors and Hsp104 colocalize in MitoStores

To verify that mitochondrial precursors and Hsp104 indeed colocalize in cytosolic granules, we fused the three mitochondrial Hsp104 clients Aim17, Pdb1 and Mam33 to the red fluorescent protein (RFP) mCherry and coexpressed them with Hsp104-GFP (Fig. 5A, B, S5C). Upon clogger induction, the RFP signal formed defined punctae that colocalized with Hsp104-GFP. Whereas Aim17-RFP binding to Hsp104-GFP was seen even without clogger induction, Pdb1 and Mam33 were only found in cytosolic granules when import was slowed down by clogger expression. Thus, the association of individual proteins with cytosolic MitoStores likely depends on the individual protein and the prevailing cellular conditions. It should be noted that Hsp104-GFP-containing granules were frequently found in yeast cells even in the absence of the clogger or uncouplers as long as cells were grown on non-fermentable carbon sources which induce the expression of mitochondrial proteins (Fig. 5C).

**Fig. 5.**
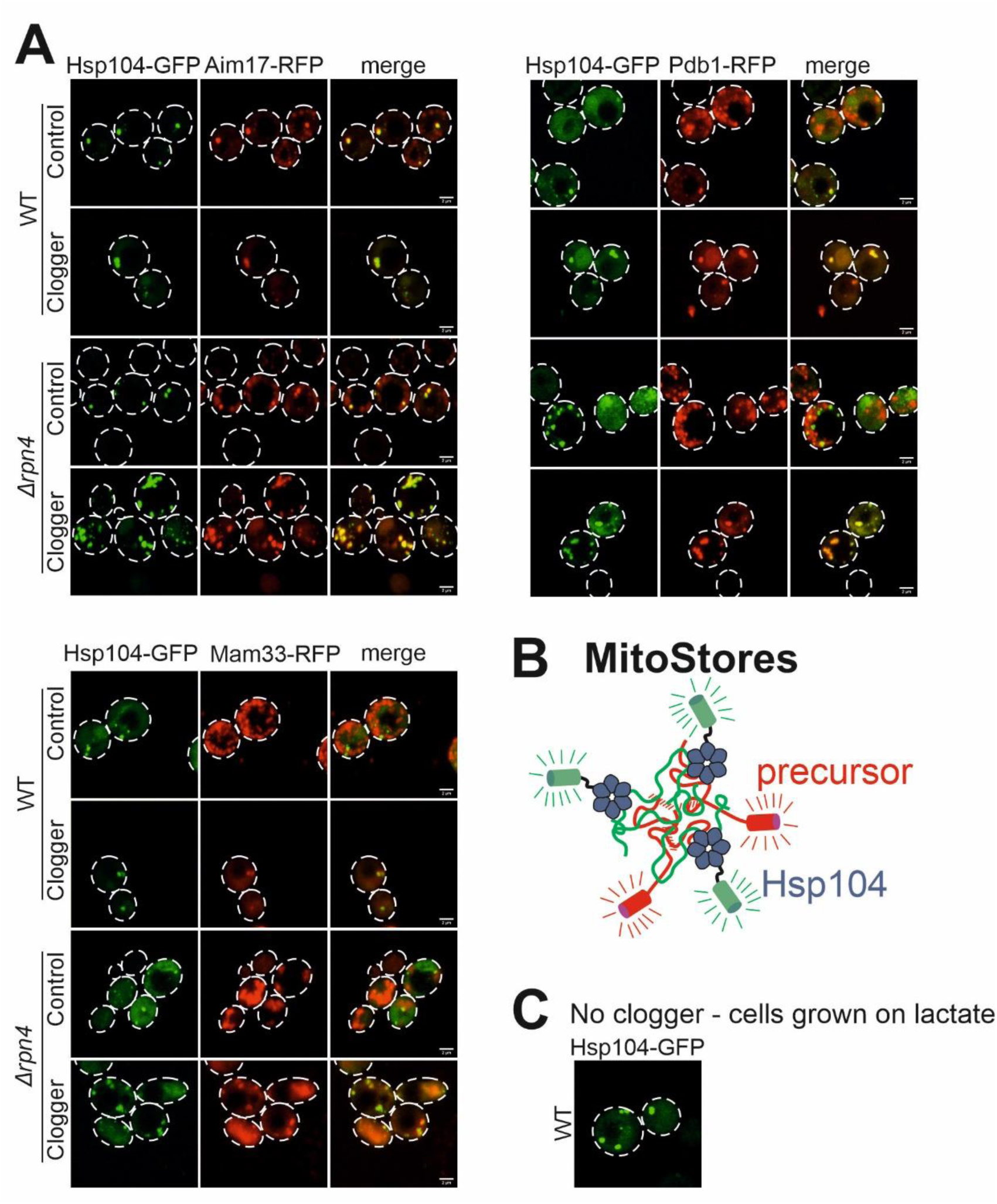
Mitochondrial proteins colocalize with Hsp104 in MitoStores. (**A**) The mitochondrial proteins Pdb1, Mam33 and Aim17 were co-expressed with clogger as fusions with the red fluorescent protein RFP (mCherry). Fluorescence microscopic images of the indicated mitochondrial proteins visualized 4.5 h after induction of the clogger or cytosolic DHFR for control. For quantification of colocalization see Fig. S5C. (**B**) Mitochondrial proteins colocalize with Hsp104-GFP in MitoStores. (**C**) Wild type cells expressing Hsp104-GFP were grown on lactate medium to log phase. Please note that aggregates are visible in many cells even in the absence of clogger expression.

### MitoStores facilitate the productive import into mitochondria

Next, we asked whether cells could resolve MitoStores once the stress conditions are relieved. To this end, we shifted clogger-inducing cells after 4.5 h continuous growth on galactose (i.e. inducting conditions) to lactate medium lacking galactose for 4 h. Once clogger induction was stopped, Hsp104 was again evenly distributed throughout the cytosol and Pdb1-RFP and Aim17-RFP reached the punctate distribution that characterizes mitochondria (Fig. 6A, S5A). To verify that mitochondrial proteins that are synthesized during the clogger-mediated import inhibition, reach the mitochondria once the block is relieved, we developed an assay relying on a split-GFP approach. To this end, we constitutively expressed the first 10 beta sheets of superfolder GFP in mitochondria (Oxa1-GFP^1-10^) and co-expressed the 11th beta sheet of GFP fused to Aim17 under a galactose-regulatable promoter. Thus, this Aim17-GFP^11^ fusion protein was co-expressed with the clogger for 4.5 h, before galactose was removed and cells were further incubated for 4 h. As shown in Fig. 6B, clogger expression reduced the access of this reporter into mitochondria; nevertheless, considerable amounts of the protein were still imported after the chase reaction (about 50% of the control sample without the clogger). Interestingly, in the absence of Rpn4, considerably higher levels were imported into the mitochondria indicating that Aim17 is indeed stabilized in the cytosol in an import-competent fashion, particularly if proteasome upregulation is prevented.

**Fig. 6.**
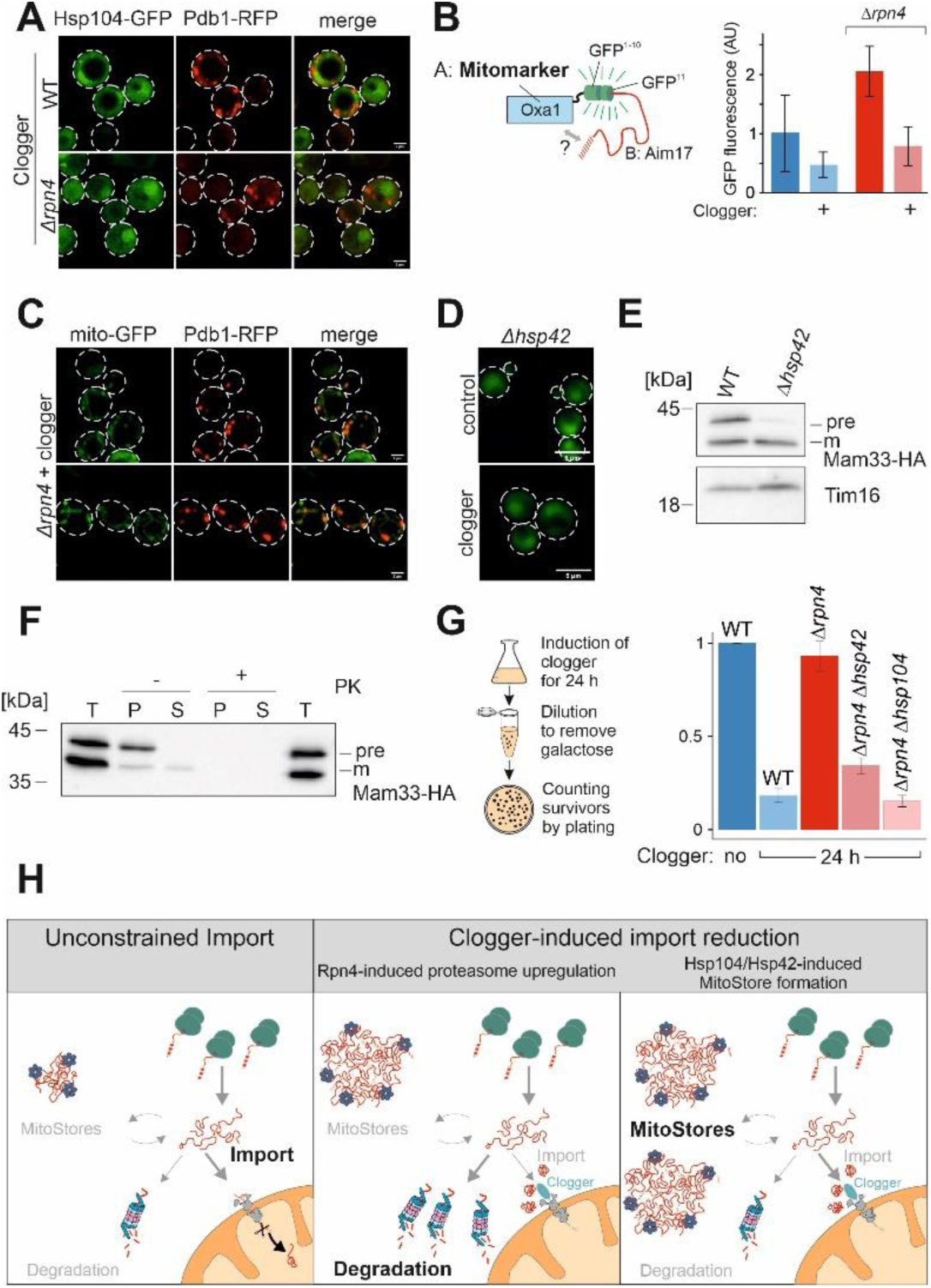
MitoStores form in proximity to mitochondria. **(A)** After 4.5 h of galactose-induced expression of clogger and Pdb1-RFP, cells were incubated for 4 h in the absence of galactose. Whereas Hsp104-GFP was distributed throughout the cytosol after this chase period, Pdb1-RFP showed a distribution pattern characteristic for mitochondrial proteins. (**B**) To verify that stored proteins are chased into the mitochondrial matrix, Aim17-GFP11 was coexpressed with clogger for 4.5 h in cells containing mitochondria with Oxa1-GFP1-10. Cells were washed in galactose-free buffer and further incubated for 4 h before fluorescence was measured to quantify the matrix-localized Aim17 proteins. (**C**) Clogger was expressed for 4.5 h before colocalization of MitoStores (visualized by Pdb1-RFP) and mitochondria (stained with matrix-residing mito-GFP) was assessed. (**D**) Clogger was expressed for 4.5 h in *Δhsp42* cells before Hsp104-GFP was visualized. Please note that in the absence of Hsp42 no aggregates were detected. (**E**) Clogger and Mam33-HA were co-expressed for 4.5 h in wild type and *Δhsp42* cells. Mitochondria were isolated and subjected to Western blotting to detect mature (m) and precursor (pre) forms of Mam33-HA. The matrix protein Tim16 served as loading control. (**F**) The mitochondria used for D were treated with proteinase K (PK) to remove surface-exposed proteins. Mitochondrial membranes were lysed with NP-40 and soluble (S) and aggregated (P) proteins were separated by centrifugation. T, total. (**G**) To measure the toxicity of clogger expression, clogger proteins were induced for 24 h in the mutants indicated. Aliquots were removed and the number of living cells were assessed by a plating assay. The resistance of *Δrpn4* to clogger expression depends on Hsp104 and Hsp42. (**H**) Schematic representation of cytosolic MitoStores.

Next, we asked where MitoStores reside in cells. Double staining of mitochondrial GFP and Pdb1-RFP indicated that MitoStores are in proximity to mitochondria (Fig. 6C, S5D). We even observed non-imported Mam33-HA precursor co-isolating with purified wild type but not *Δhsp42* mitochondria from clogger-stressed cells (Fig. 6E). This precursor was protease-accessible and found in high-speed pellets after solubilizing mitochondrial membranes with detergent, indicating that MitoStores remain associated with the mitochondrial surface even during cellular fractionation procedures (Fig. 6E). Since Hsp42 was crucial for the formation of MitoStores (Fig. 6D) we tested whether deletion of Hsp42 suppressed the clogger resistance observed in *Δrpn4* mutants (Fig. 6F). Indeed, *Δrpn4* cells required Hsp42 and Hsp104 to survive clogger stress (Fig. 6G). Thus, Hsp42 and Hsp104 protect against clogger stress in the absence of Rpn4 which is reminiscent of their role in heat stress (Fig. S5E, S5F). In summary, we describe here that the cytosol of yeast cells has the capacity to transiently store mitochondrial matrix proteins in Hsp42-induced and Hsp104-bound structures which we name MitoStores (Fig. 6H). These structures reduce the toxic effects arising from non-imported mitochondrial precursor proteins.

## Discussion

To our surprise we observed that yeast cells lacking Rpn4 can tolerate the accumulation of mitochondrial precursors in their cytosol even better than wild type cells. Instead of degrading these precursors, these cells efficiently and selectively package precursors into cytosolic granules that are bound by the disaggregase Hsp104. Hsp42 and Hsp104 are upregulated in Drpn4 cells even before clogger expression explaining the increased resistance against mitoprotein-induced stress of these cells.

These MitoStores resemble stress granules or Q-bodies in their overall appearance (Eisele et al., 2021; Escusa-Toret et al., 2013; Hill et al., 2017; Roth and Balch, 2013; Samant et al., 2018), however, they largely contain matrix-destined mitochondrial precursor proteins as well as precursors of secretory proteins. We observed that MitoStores transiently accumulate during phases at which the synthesis of mitochondrial proteins exceeds the capacity of the mitochondrial import system. This is most extreme upon clogger-mediated competitive inhibition of protein import, but also apparent when cells are grown on non-fermentative carbon sources such as lactate or glycerol as well as at late stages on glucose media where cells undergo a diauxic shift to switch to ethanol consumption (Di Bartolomeo et al., 2020).

The specific upregulation of Hsp42 and Hsp104 in Rpn4-deficient cells indicates that MitoStore formation and proteasome-mediated degradation are two alternative strategies to deal with the accumulation of mitochondrial precursors in the cytosol (Fig. 6G). The balance of both processes is not clear but the immediate Rpn4-mediated induction of the proteasome system (Boos et al., 2019) indicates that cytosolic precursors are under tight proteolytic control. The proteasome plays a direct and crucial role in the removal of stalled import intermediates from the mitochondrial outer membrane translocase or from other cellular non-productive binding sites (Basch et al., 2020; Itakura et al., 2016; Mårtensson et al., 2019; Mohanraj et al., 2019; Murschall et al., 2020; Shakya et al., 2021; Weidberg and Amon, 2018). Hence, proteasomal upregulation appears to be the primary and immediate cellular response to mitoprotein-induced stress conditions. Nevertheless, cells actively promote the formation of MitoStores in the cytosol employing small heat shock proteins such as Hsp26 and Hsp42 as nucleation sites (Haslbeck et al., 2020; Mogk and Bukau, 2017), presumably to sequester precursors to reduce their toxicity. Interestingly, MitoStores apparently accommodate a specific subset of proteins, in particular proteins with N-terminal presequences. The proteins that are typically used in the field to detect import defects by detection of precursor proteins in Western Blots, such as Rip1, Mdj1, Hsp60 or Sod2, were all found highly enriched in MitoStores (Gallas et al., 2006; Mårtensson et al., 2019; Nowicka et al., 2021; Poveda-Huertes et al., 2020; Sinha et al., 2010). Strikingly, members of the carrier family were not detected in MitoStores even though this group of inner membrane proteins is very abundant (Morgenstern et al., 2017) and exhibits a highly toxic potential (Wang and Chen, 2015; Xiao et al., 2021). Apparently, the import of carrier proteins deviates from that of presequence-containing precursors at early stages of the cytosolic targeting. Consistently, previous studies reported that these two groups employ different factors of the cytosolic chaperone system during their targeting to mitochondria (Backes et al., 2021; Hansen et al., 2018; Komiya et al., 1997; Opalinski et al., 2018). These variations in the pre-mitochondrial life of mitochondrial proteins might explain the differences between the mitoprotein-induced stress reactions towards accumulating presequence-containing proteins (typically referred to as UPRam) and accumulating carriers (referred to as mitochondrial precursor over-accumulation stress, mPOS)(Wang and Chen, 2015; Wrobel et al., 2015). The accumulation of carriers is apparently much more toxic, potentially because they are not sequestered in MitoStores (Hoshino et al., 2019; Pouikli et al., 2021; Wang et al., 2022) but rather interfere with non-mitochondrial membranes (Xiao et al., 2021).

We observed that MitoStores are transient in nature and dissolve once the mitoprotein-inducing stress conditions are ameliorated. This is different to what was reported for IPODs or aggresomes which are toxic aggregates that form for example from insoluble polyQ proteins (Hill et al., 2014; Schlagowski et al., 2021; Spokoini et al., 2012; Tyedmers et al., 2010).

Whereas IPOD formation uses the yeast prion Rnq1 for nucleation, we observed no relevance of Rnq1 for the formation of MitoStores (Fig. S5B). We therefore favor the idea that MitoStore are benign structures which safeguard cellular proteostasis during overload of the mitochondrial import system or metabolic remodeling.

Our observation that mitochondrial precursor proteins can be selectively incorporated into dedicated storage granules in the cytosol adds another step into the cascade of reactions that mediate the targeting and import of mitochondrial proteins. Further studies will have to elucidate the relevance of cytosolic factors and of the ubiquitination machinery to control the stress-induced formation and post-stress resolution of MitoStores and to characterize their physical and functional association with proteins of the mitochondrial surface.

## Acknowledgements

We thank Vera Nehr, Sabine Knaus and Johannes Ebding for technical assistance. We thank Christian Koch and Carina Groh for the help with data analysis. We thank Per Haberkant and Mikhail Savitski for help with establishing the TMT/mass spectrometry workflow and Dieter Wolf, Martin van der Laan, Günter Kramer, Thomas Becker and Dejana Mokranjac for reagents. This study was supported by the Deutsche Forschungsgemeinschaft (GRK2737 *STRESSistance* to J.M.H., Z.S. J.P. and HE2803/10-1 to J.M.H.), the Joachim Herz Stiftung (to F.B.), the Damon Runyon Cancer Research Foundation (DRG-2461-22 to F.B.) and the Forschungsinitiative Rheinland-Pfalz BioComp (to J.M.H., F.B., Z.S. and J.P.).

## Author Contributions

L.K. designed, cloned and verified the constructs and strains; F.B. and J.M.H. conceived the project; M.R. and Z.S. carried out the mass spectrometry-based proteomics; F.B., M.R. and L.K. performed the bioinformatical analysis of the mass spectrometry data; L.K. carried out the biochemical experiments to test the relevance of Rpn4 in the context of the mitoprotein-induced stress response; L.K., N.D. and J.P. carried out the fluorescence microscopy experiments and carried out the image analysis; L.K., N.D. and J.M.H. analyzed the data; J.M.H. and L.K. wrote the manuscript with the help and input of all authors.

## Declaration of Interests

The authors declare no competing financial interests.

---

**Fig S1.**
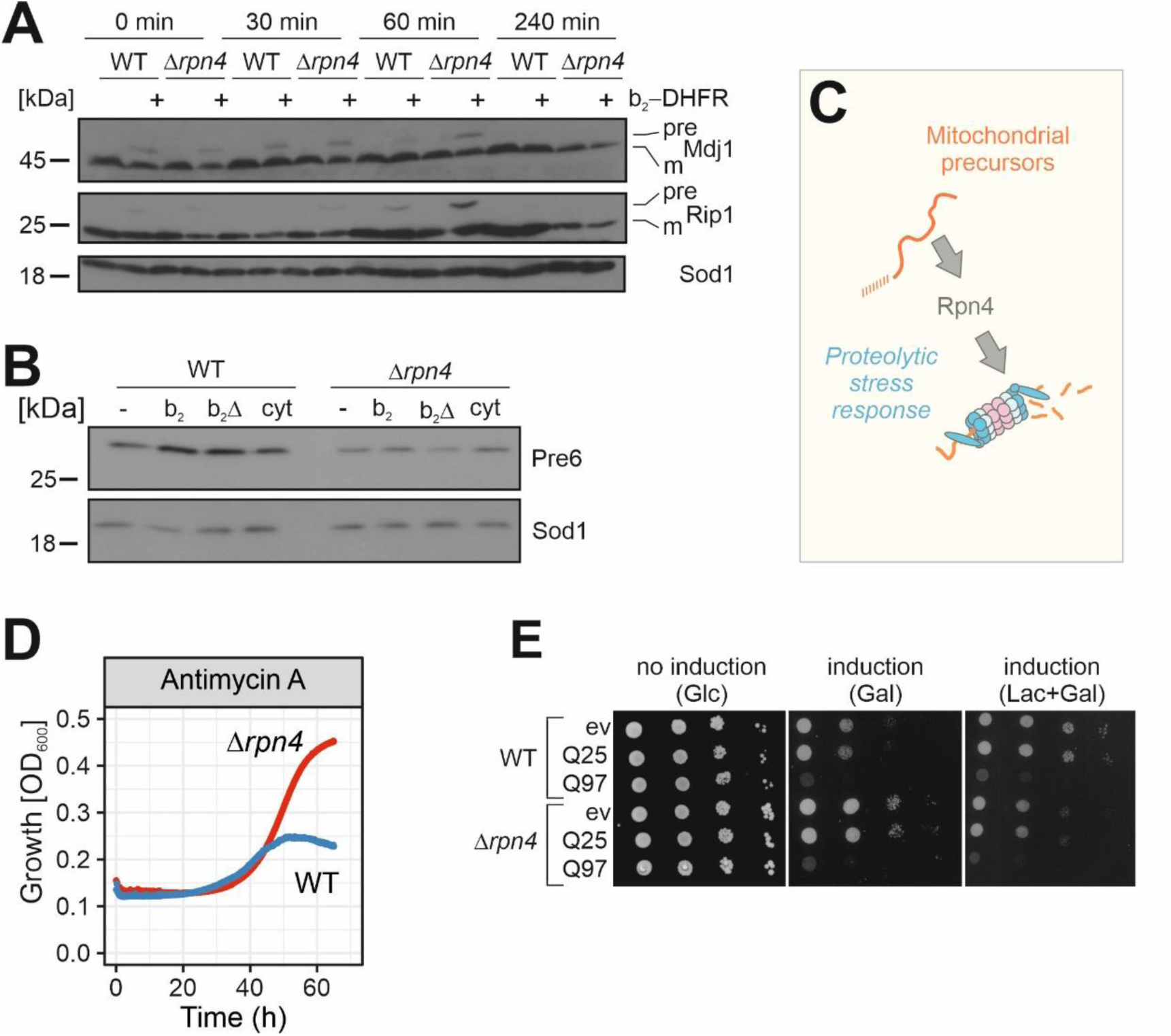
Rpn4-dependent proteasome induction. **(A)** After 4.5 h of clogger expression in WT and *Δrpn4* cells, medium was exchanged for a non-inducing lactate medium. Precursor (pre) and mature (m) forms of the mitochondrial proteins Mdj1 and Rip1 were visualized by Western blotting. Sod1 was used as a loading control. **(B)** After 4.5 h expression of clogger and control, protein levels of the proteasome subunit Pre6 were visualized by Western Blotting. Please note, that in *Δrpn4* cells, clogger expression does not lead to Pre6 induction. Sod1 served as loading control. **(C)** Schematic representation of Rpn4-dependent proteasome induction after mitoprotein-induced stress. **(D)** Cells of the indicated strains were grown to log phase and diluted in lactate medium to 0.1 OD. After addition of 100 µg/ml antimycin A, growth was monitored upon constant agitation. **(E)** Yeast cells expressing a fusion protein of the N-terminal region of human huntingtin with different lengths of glutamine residues (Q25-GFP and Q97-GFP, respectively) (Schlagowski et al., 2021) and GFP under control of the galactose-inducible promoter, or a control for comparison were grown to mid-log phase on non-inducing glucose medium. Ten-fold serial dilutions were dropped on glucose (no induction), galactose (induction) or lactate with 0.5% galactose medium (induction).

**Fig. S2.**
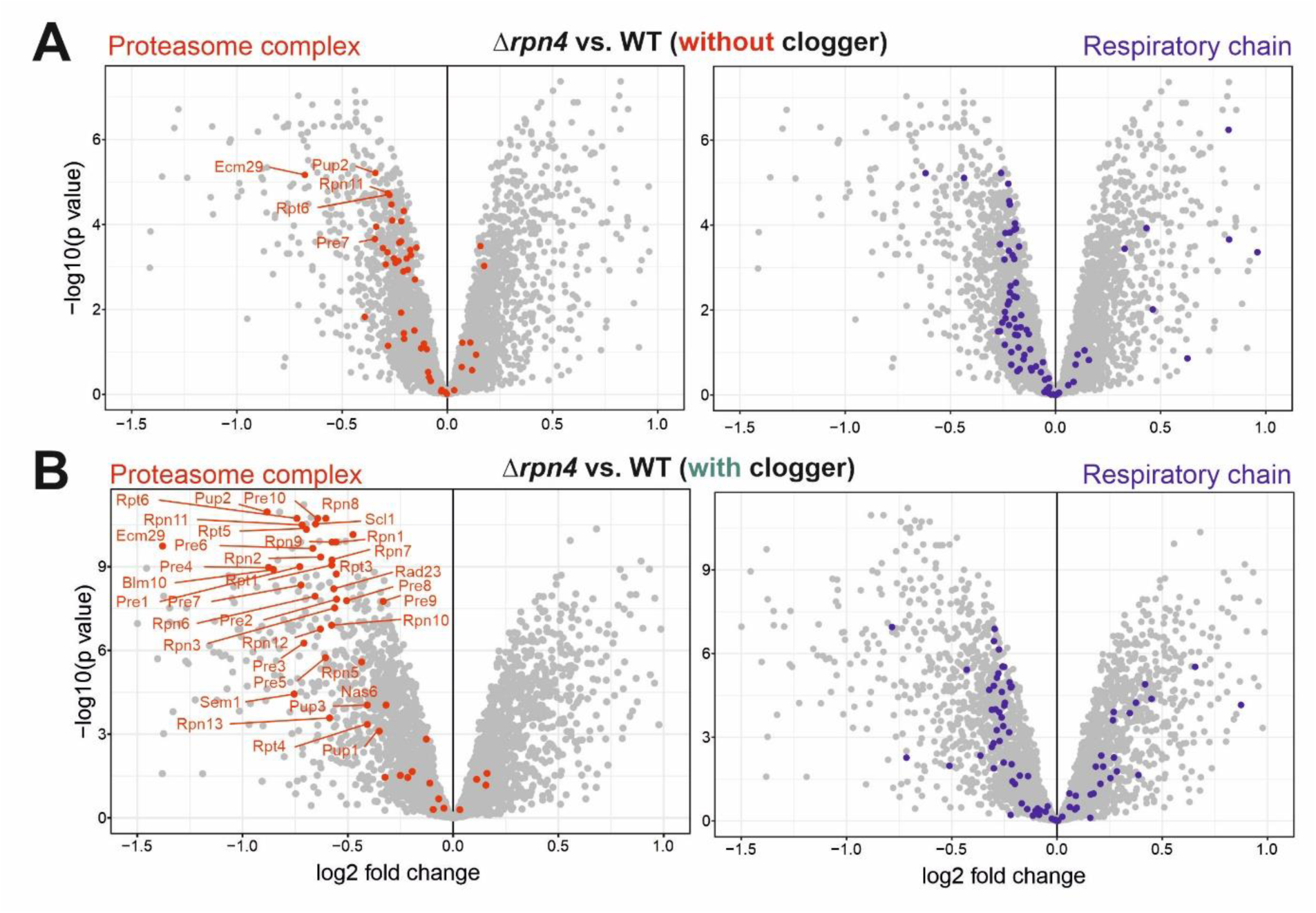
Rpn4 deletion leads to a reduction in proteasomal and respiratory proteins. **(A)** Comparison of the proteome of wild type and *Δrpn4* cells before and after clogger expression for 6 h. Positions of proteins of the proteasome and the respiratory chain systems are indicated in red and blue, respectively (Boos et al., 2019; Morgenstern et al., 2017). Please note especially the proteasomal proteins show a lower abundance in *Δrpn4* cells.

**Fig. S3.**
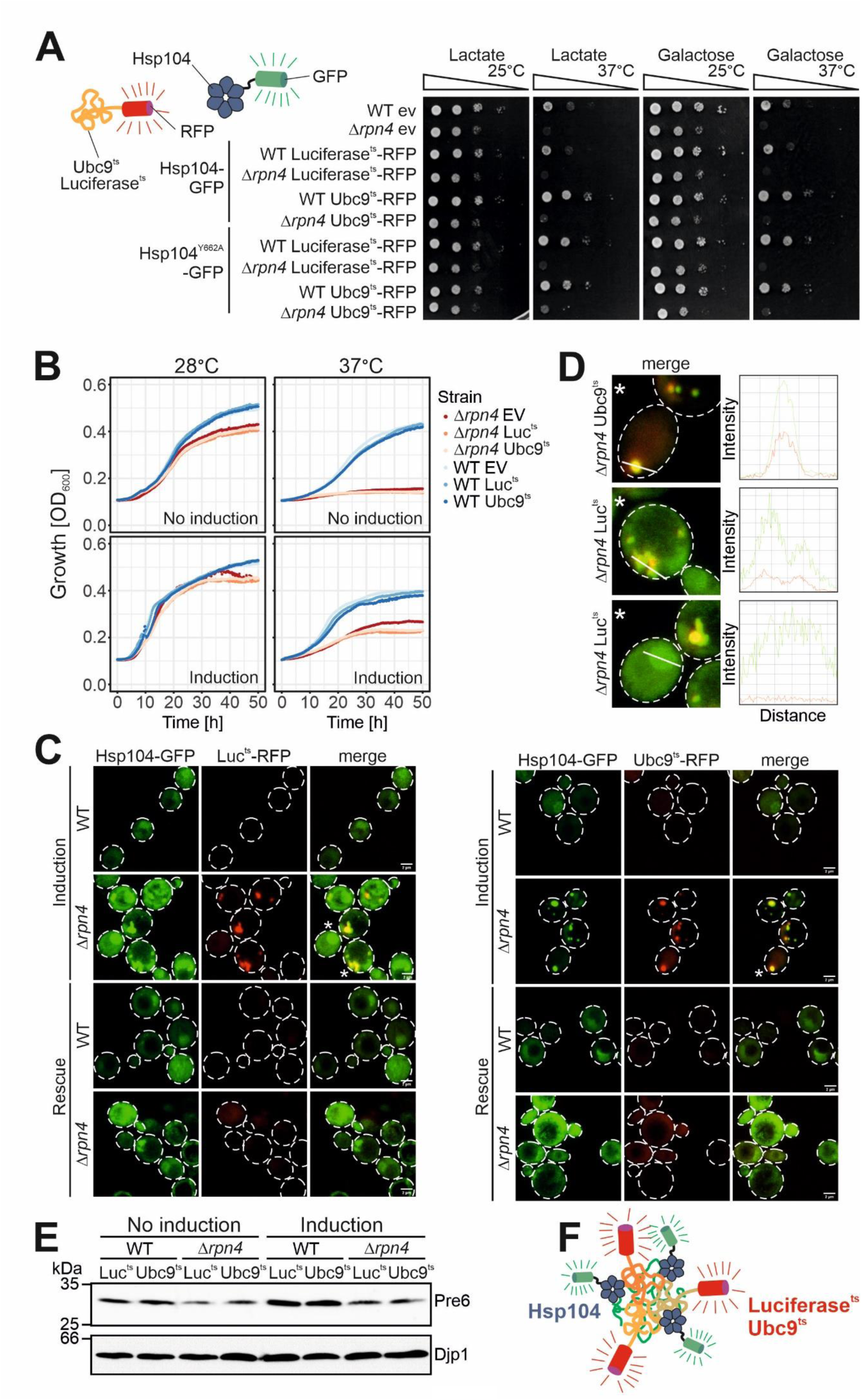
Protein aggregation in the yeast cytosol can be efficiently measured by visualization of the distribution of Hsp104-GFP. **(A)** Yeast cells expressing the temperature-sensitive, aggregation-prone proteins luciferase^ts^-RFP (Luc^ts^-RFP) and Ubc9^ts^-RFP under galactose-inducible promoters together with constitutively expressed Hsp104-GFP or Hsp104^Y662A^-GFP (Lum et al., 2004) were grown to mid-log phase on lactate medium. Ten-fold serial dilutions were dropped on lactate (no induction) or galactose medium (induction) and incubated at 25°C or 37°C as indicated. **(B)** Cells of the indicated strains were grown to log phase and diluted in lactate (no induction) or galactose (induction) to 0.1 OD. Growth was monitored upon constant agitation at 28°C and 37°C, respectively. **(C)** After 2 h of Luc^ts^-RFP or Ubc9^ts^-RFP expression together with constitutive Hsp104-GFP expression at 37°C (induction), cells were incubated for 4 h in the absence of galactose at 30°C (rescue). **(D)** Fluorescence intensity profiles of cells marked with an asterisk in (C). The measured area is indicated with a white line. **(E)** After 4 h growth on lactate medium (no induction) or galactose medium (induction), protein level of the proteasomal protein Pre6 was visualized by Western Blotting. Djp1 was used as a loading control. **(F)** Schematic representation of the interaction of aggregation-prone proteins with Hsp104.

**Fig. S4.**
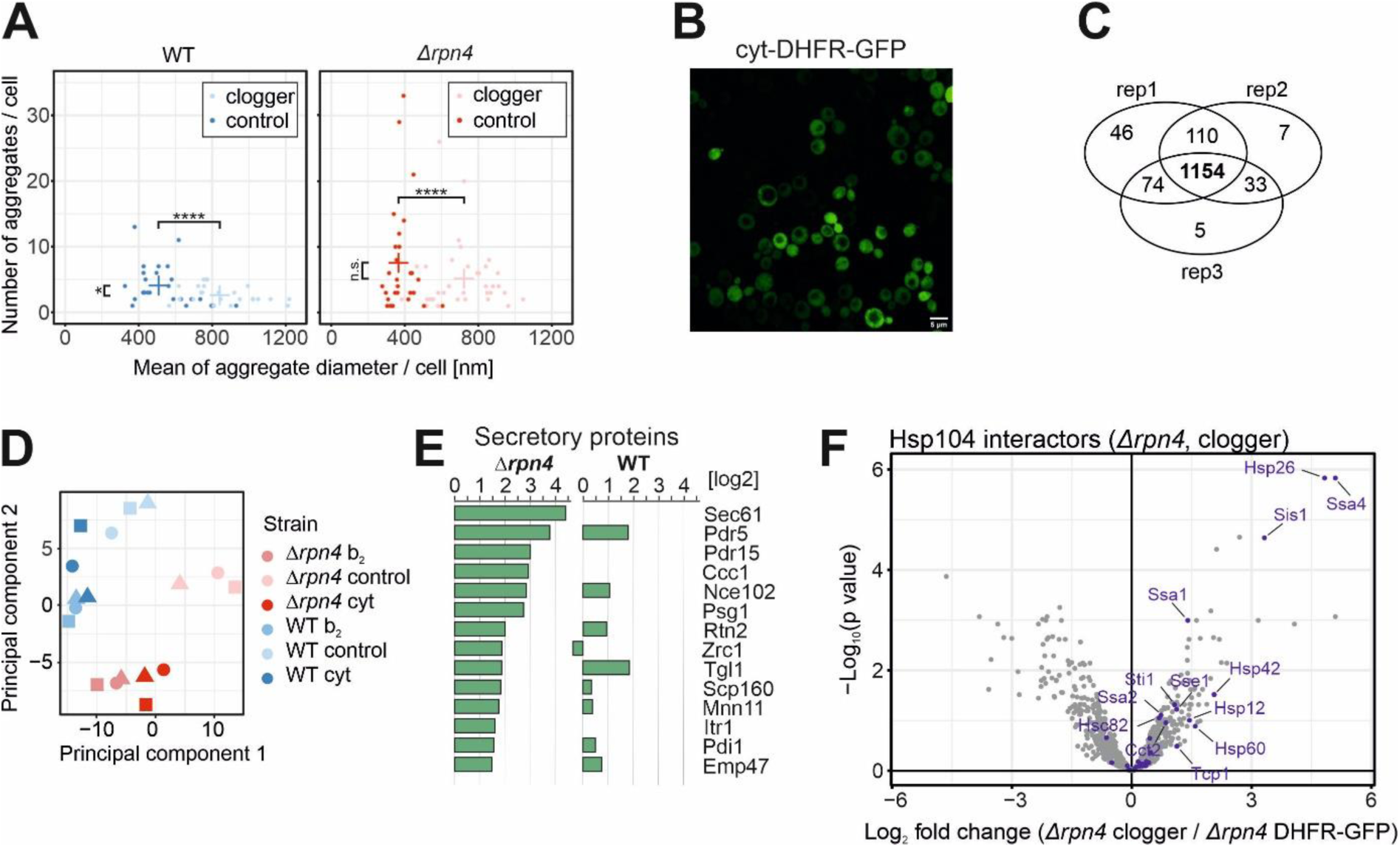
Deletion of Rpn4 remodels the cytosolic chaperone network. **(A)** Quantification of aggregate formation in WT and *Δrpn4* cells expressing either the clogger or a cytosolic DHFR as a control for 4.5 h. Plotted are the mean diameters of the individual aggregates in nm, as well as the number of aggregates per cell. **(B)** After expressing the cytosolic DHFR-GFP construct for 4.5 h in WT cells, the expression was visualized by confocal microscopy. **(C)** Identification overview of the mass spectrometry. 1,154 proteins were measured in all samples of the three replicates. **(D)** Principal component analysis. Rpn4 deletion and clogger expression caused specific changes in the proteome. The different shapes of the individual data points indicate the three biological replicates **(E)** Purified proteins of the secretory pathway interacting with Hsp104-GFP in *Δrpn4* expressing the clogger normalized to the control. **(F)** Hsp104 interactors in the presence of clogger relative to interactors of DHFR-GFP. Chaperones are shown in purple.

**Fig. S5.**
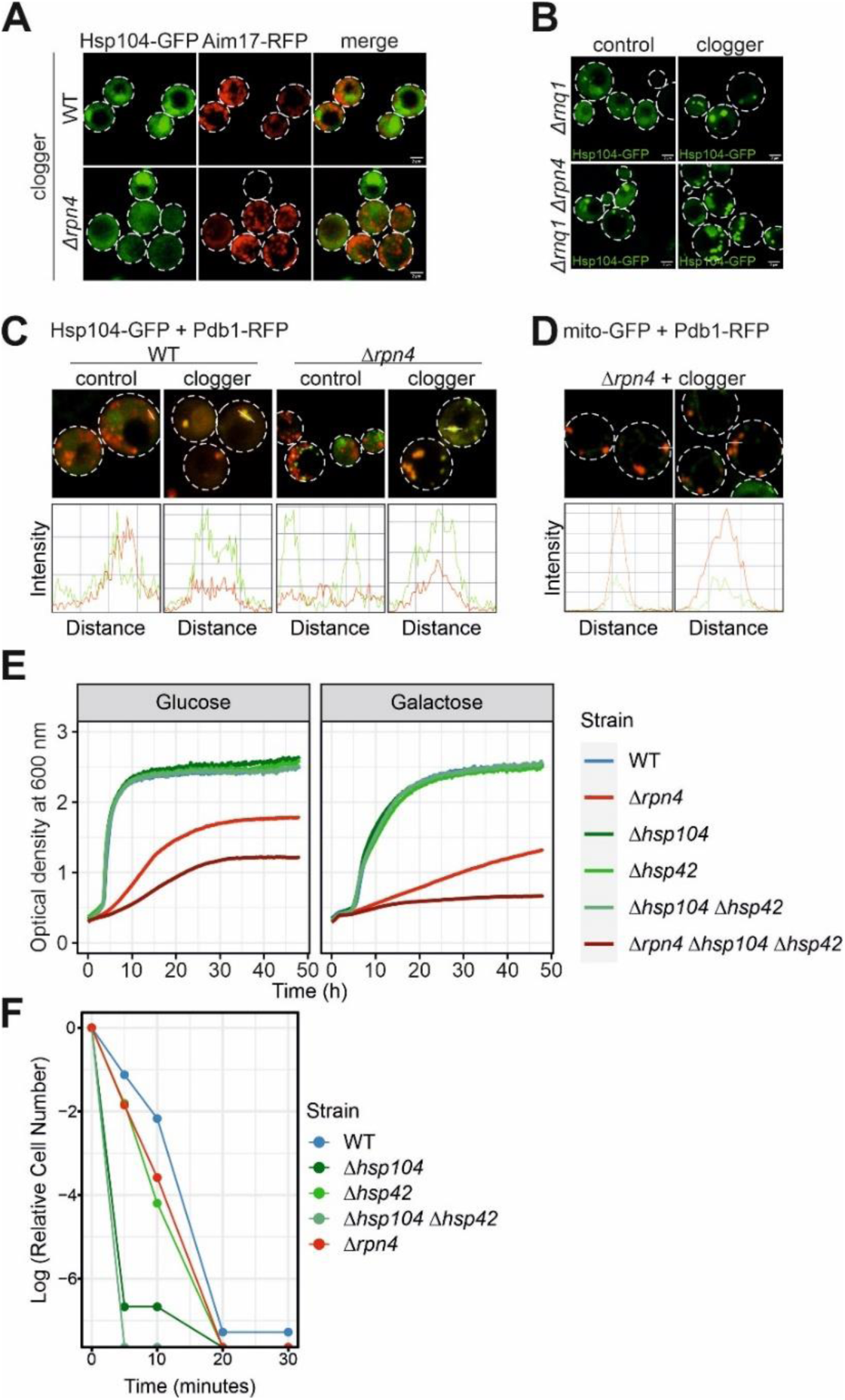
MitoStore formation depends on the Hsp42 / Hsp104 chaperone system. **(A)** After 4.5 h of clogger and Aim17-RFP expression, WT and *Δrpn4* cells were incubated for 4 h in the absence of galactose. Whereas Hsp104-GFP was distributed throughout the cytosol after this chase period, Aim17-RFP showed a distribution pattern characteristic for mitochondrial proteins. **(B)** Microscopy of *Δrnq1* and *Δrpn4Δrnq1* cells expressing Hsp104-GFP constitutively and the clogger or cytosolic control for 4.5 h. Please note that the formation of MitoStores did not depend on the yeast prion protein Rnq1. **(C)** Quantification of the colocalization of Hsp104-GFP with Pdb1-RFP after clogger expression for 4.5 h. **(D)** Quantification of the colocalization of mito-GFP with Pdb1-RFP after clogger expression for 4.5 h in the *Δrpn4* mutant. **(E)** Cells of the indicated strains were grown to log phase and diluted in glucose or galactose medium to 0.1 OD. Growth was monitored upon constant agitation at 30°C or 37°C respectively. **(F)** Cells were grown in lactate to mid log phase before 50°C heat stress was performed for the indicated time points. After each time aliquots were removed, and the number of living cells were assessed by a plating assay.

## METHOD DETAILS

### Strains and growth conditions

The yeast strains and plasmids used in this study are described in detail in Supplementary Table S3 and S4, respectively. Unless specified, all strains were derived from YPH499*Δarg4* (MATa *ura3 lys2 ade2 trp1 his3 leu2*).

The strains were grown at 30°C either in yeast complete medium (YP) containing 1% (w/v) yeast extract, 2% (w/v) peptone and 2% (w/v) of the respective carbon source or in minimal synthetic respiratory medium containing 0.67% (w/v) yeast nitrogen base and 2% lactate as carbon source. To induce the clogger from the *GAL1* promoter, 0.5% galactose was added. The temperature-sensitive strains and its corresponding wildtype were grown at 25°C and the phenotype was induced with growth at 37°C.

### Growth assays and viability tests

For spot analysis, the respective yeast strains were grown in liquid rich or synthetic media. Total yeast cells equivalent to 0.5 OD_600_ were harvested at exponential phase. The cells were washed in sterile water and subjected to ten-fold serial dilutions. From each dilution, 3 µl were spotted on the respective media followed by incubation at 30°C or 37°C. Pictures were taken after different days of the growth.

Growth curves were performed in a 96 well plate, using the automated ELx808™ Absorbance Microplate Reader (BioTek®). The growth curves started at 0.1 OD_600_ and the OD_600_ was measured every 10 min for 72 h at 30°C or 37°C. The mean of technical triplicates was calculated and plotted in R.

For viability assays, yeast cells were pre-grown in lactate medium at 30°. At mid-log phase (OD_600_ 0.6-0.8), either clogger expression was induced with 0.5% galactose or the cells were exposed to 50°C for the indicated time points. After each timepoint, 100 µl of 0.001 OD_600_ cell suspension were equally plated on lactate plates. The plates were incubated at 30°C for 3 days to assess the number of colonies which survived the mitoprotein-induced or respectively the heat stress.

### Antibodies

The antibodies against Sod1, GFP and Djp1 were raised in rabbits using recombinant purified proteins. The antibody against Pre6 was kindly gifted by Dieter Wolf (University of Stuttgart, Germany). The antibody against mouse DHFR was kindly provided by Martin van der Laan (Saarland University, Germany). The antibodies against Rip1 and Mdj1 were kindly gifted by Thomas Becker (University of Freiburg, Germany). The antibody against Ssa1 was kindly provided by Günter Kramer (ZMBH Heidelberg, Germany). The antibody against Tim16 was kindly gifted by Dejana Mokranjac (LMU München, Germany). The horseradish-peroxidase coupled HA antibody was ordered from Roche (Anti-HA-Peroxidase, High Affinity 3F10), #12 013 819 001). The secondary antibodies were ordered from Biorad (Goat Anti-Rabbit IgG (H+L)-HRP Conjugate #172-1019). Antibodies were diluted in 5% (w/v) nonfat dry milk-TBS (Roth T145.2) with the following dilutions: anti-Sod1 1:1,000, anti-Rip1 1:750, anti-Mdj1 1:125, anti-Ssa1 1:10,000, anti-Rabbit 1:10,000

### YFP reporter assays

The PACE-YFP reporter gene was integrated into the *LEU2* locus of the yeast genome. Cells were grown to mid log phase (OD 0.6 – 0.8) under non-inducing conditions and shifted to inducing conditions by addition of 0.5% galactose for 4.5 hours. 4 OD_600_ of cells were harvested by centrifugation (20,000g, 3 min, RT) and resuspended in 400 µl H_2_O. 100 µl of cell suspension were and transferred to flat-bottomed black 96-well imaging plates (BD Falcon, Heidelberg, Germany) in technical triplicates. Cells were sedimented by gentle spinning (30 g, 5 min, RT) and fluorescence (excitation 497 nm, emission 540 nm) was measured using a ClarioStar Fluorescence Platereader (BMG-Labtech, Offenburg, Germany). The corresponding wildtype strain not expressing YFP was used for background subtraction of autofluorescence.

Fluorescence intensities were normalized to the value obtained from the wildtype empty vector control in each of three independent biological replicates. Statistical significance was assessed using a paired one-tailed Student’s t-test where a p value of below 0.05 was interpreted as significant difference.

### Cell lysates

For whole cell lysates, yeast strains were cultivated in selective lactate media to mid-log phase. Clogger expression was induced by the addition of 0.5% galactose. For recovery assays the medium was exchanged for a non-inducing lactate medium. After the indicated timepoints 4 OD_600_ were harvested by centrifugation (17,000 g, 2 min). Cells were washed with water and resuspended in reducing loading buffer. Cells were transferred to screw cap tubes containing 1 mm glass beads. Cell lysis was performed using a FastPrep-24 5G homogenizer (MP Biomedicals, Heidelberg, Germany) with 3 cycles of 30 s, speed 8.0 m/s, 120 s breaks, glass beads). Lysates were boiled at 96°C for 5 min, centrifuged (17,000 g, 2 min) and stored at - 20°C until further use. The equal amount of OD_600_ was loaded on an SDS-gel.

### Isolation of mitochondria

For the isolation of mitochondria cells were grown in selective lactate media. Cells were harvested (4,000 rpm, JA10 Beckmann rotor, 5 min, RT) in the exponential phase. After a washing step, cells were treated 10 min with 2 ml per g wet weight MP1 buffer (10 mM Tris pH unadjusted and 100 mM DTT) at 30°C. After washing with 1.2 M sorbitol, yeast cells were resuspended in 6.7 ml per g wet weight MP2 buffer (20 mM KPi buffer pH 7.4, 1.2 M sorbitol, 3 mg per g wet weight zymolyase from Seikagaku Biobusiness) and incubated for 1 h at 30°C. Spheroplasts were collected via centrifugation at 4°C and resuspended in ice cold homogenization buffer (13.4 ml/g wet weight) (10 mM Tris pH 7.4, 1 mM EDTA pH 8, 0.2% fatty acids free bovine serum albumin (BSA), 1 4 mM PMSF, 0.6 M sorbitol). Spheroplasts were disrupted by 10 strokes with a cooled glass potter. Cell debris was removed via centrifugation at 3,300 rpm in a JA10 Beckmann rotor. The supernatant was centrifuged for 12 min at 10,000 rpm to collect mitochondria. Mitochondria were resuspended in 10 ml of ice cold SH-buffer (0.6 M sorbitol, 20 mM Hepes pH 7.4) and centrifuged again at 4,000 rpm in a JA25.50 Beckmann rotor to remove residual cell debris. To harvest the mitochondria the supernatant was again centrifuged for 12 min at 12,000 rpm. The amount of mitochondria was determined using the Bradford assay.

### Sample preparation and mass spectrometric identification of proteins

For the quantitative comparison of proteomes of *Δrpn4* and WT cells expressing the clogger, expression of b_2_-DHFR was induced for 6 h and compared to control cells expressing cytosolic DHFR. 10 OD_600_ of cells were harvested at each time point by centrifugation (17,000 g, 3 min, 2°C), washed with prechilled water, snap-frozen in liquid nitrogen and stored at -80°C. Cells lysates were prepared in lysis buffer (50 mM Tris pH 7.5, 2% (w/v) SDS, Tablets mini EDTA-free protease inhibitor (Roche)) using a FastPrep-24 5G homogenizer (MP Biomedicals, Heidelberg, Germany) with 3 cycles of 30 s, speed 8.0 m/s, 120 s breaks, glass beads). Lysates were boiled for 5 min at 96°C and centrifuged (17,000 g, 3 min, 2°C). Protein concentrations were determined using the Pierce BCA Protein Assay (Thermo Scientific, #23225). 20 µg of each lysate were subjected to an in-solution tryptic digest using a modified version of the Single-Pot Solid-Phase-enhanced Sample Preparation (SP3) protocol^48,49^. Here, lysates were added to Sera-Mag Beads (Thermo Scientific, #4515-2105-050250, 6515-2105-050250) in 10 µl 15% formic acid and 30 µl of ethanol. Binding of proteins was achieved by shaking for 15 min at room temperature. SDS was removed by 4 subsequent washes with 200 µl of 70% ethanol. Proteins were digested with 0.4 µg of sequencing grade modified trypsin (Promega, #V5111) in 40 µl Hepes/NaOH, pH 8.4 in the presence of 1.25 mM TCEP and 5 mM chloroacetamide (Sigma-Aldrich, #C0267) overnight at room temperature. Beads were separated, washed with 10 µl of an aqueous solution of 2% DMSO and the combined eluates were dried down. In total three biological replicates were prepared (n=3). Each replicate included samples of control and clogger expressing cells (in total 6 samples per replicate). Peptides were reconstituted in 10 µl of H_2_O and reacted with 80 µg of TMT10plex (Thermo Scientific, #90111)^22^ label reagent dissolved in 4 µl of acetonitrile for 1 h at room temperature. Excess TMT reagent was quenched by the addition of 4 µl of an aqueous solution of 5% hydroxylamine (Sigma, 438227). Peptides were mixed to achieve a 1:1 ratio across all TMT-channels. Mixed peptides were desalted on home-made StageTips containing Empore C_18_ disks (Rappsilber et al., 2007) and subjected to an SCX fractionation on StageTips into 3 fractions, followed by additional cleanup on C_18_ StageTips. fractionation under high pH condition^48^. The resulting 12 fractions were then analyzed by LC-MS/MS on a Q Exactive Plus (Thermo Scientific) as previously described^50^.

For IP mass spectrometry, cells were incubated in lactate medium containing the galactose to induce the expression of the clogger. After 6 h 10 OD_600_ cells were harvested by centrifugation (5,000 g for 5 min), washed with prechilled water, snap-frozen in liquid nitrogen and stored at -80°C. Cell lysates were prepared in 200 µl ice-cold lysis buffer (25 mM Tris-HCl pH 7.5, 50 mM KCl, 10 mM MgCl_2_, 5% (v/v) glycerol, 1% Nonidet P-40, 1mM DTT, 1mM PMSF, 1x cOmplete™ Tablets mini EDTA-free protease inhibitor (Roche), PhosSTOP phosphate inhibitor (Roche)) using a FastPrep-24 5G homogenizer (MP Biomedicals, Heidelberg, Germany) with 3 cycles of 30 s, speed 8.0 m/s, 120 s breaks, glass beads).Samples were centrifuged at 5,000 g for 5 min and transferred to precooled microtubes to get rid of cell debris. 300 µl dilution buffer (10 mM Tris-HCl pH 7.5, 150 mM NaCl, 0.5 mM EDTA, cOmplete™ Tablets mini EDTA-free protease inhibitor (Roche), 1 mM PMSF) was added. Diluted cell lysates were used for an IP with GFP-Trap magnetic agarose beads from Chromotek. The GFP-Trap beads were equilibrated by washing 25 µl of beads slurry with 500 µl ice-cold dilution buffer and subsequently centrifuged for 2 min at 2,500 g. The washing step was repeated two more times. GFP-tagged proteins (Hsp104 or DHFR respectively) were bound to GFP-Trap beads for 1h at 4°C tumbling end-over-end. Samples were centrifuged (2 min at 2,500 g) and supernatant was discarded by using a magnetic rack. Beads were washed 3x with 800 µl ice-cold wash buffer I (150 mM NaCl, 50 mM Tris-HCl pH 7.5, 5% (v/v) glycerol, 0.05% Nonidet P-40) and afterwards 2x with 500 µl ice-cold wash buffer II (150 mM NaCl, 50 mM Tris-HCl pH 7.5, 5% (v/v) glycerol). Samples for Western Blotting were eluted in 1x Laemmli and boiled for 5 min at 90°C. For elution and trypsin digestion of mass spectrometry samples, 50 µl elution buffer I was added (2 M Urea, 50 mM Tris-HCl pH 7.5, 1 mM DTT, 5 ng/µl trypsin) and incubated for 1h at RT. Then, 1 µl trypsin (15 ng/µl) was added and incubated for 10 min at RT. Samples were centrifuged (2,500 g, 2 min) and supernatants were transferred to fresh tubes. 50 µl elution buffer II (2M Urea, 50 mM Tris pH 7.5, 5mM CAA) was added. Samples were incubated ON in the dark at RT. pH of samples was adjusted to pH <2 with Tri-flouracetic acid. Desalting/reversed-Phase cleanup with 3xC18 stage tips. Samples were dried down in speed-vac and resolubilized in 9 µl buffer A (0.1 % formic acid in MS grad water) and 1 µl buffer A* (0.1 % formic acid, 0.1 % TFA in MS grad water). The samples were analyzed by LC-MS/MS on a Q Exactive HF(Thermo Scientific) as previously described (Sridharan et al., 2019).

Briefly, peptides were separated using an Easy-nLC 1200 system (Thermo Scientific) coupled to a Q Exactive HF mass spectrometer via a Nanospray-Flex ion source. The analytical column (50 cm, 75 µm inner diameter (NewObjective) packed in-house with C18 resin ReproSilPur 120, 1.9 µm diameter Dr. Maisch) was operated at a constant flow rate of 250 nl/min. For TMT labeled samples, a 3 h gradient was used to elute peptides (Solvent A: aqueous 0.1% formic acid; Solvent B: 80 % acetonitrile, 0.1% formic acid). Peptides were analyzed in positive ion mode applying with a spray voltage of 2.3 kV and a capillary temperature of 250°C. MS spectra with a mass range of 375–1.400 m/z were acquired in profile mode using a resolution of 120.000 [maximum fill time of 80 ms or a maximum of 3e6 ions (automatic gain control, AGC)]. Fragmentation was triggered for the top 15 peaks with charge 2–8 on the MS scan (data-dependent acquisition) with a 30 s dynamic exclusion window (normalized collision energy was 32). Precursors were isolated with a 0.7 m/z window and MS/MS spectra were acquired in profile mode with a resolution of 60,000 (maximum fill time of 100 ms, AGC target of 1e5 ions, fixed first mass 100 m/z).

For label-free IP-MS samples, shorter gradients (90 minutes) were used to elute peptides (Solvent A: aqueous 0.1% formic acid; Solvent B: 80 % acetonitrile, 0.1% formic acid). MS spectra with a mass range of 300–1.650 m/z were acquired in profile mode using a resolution of 60.000 [maximum fill time of 20 ms or a maximum of 3e6 ions (automatic gain control, AGC)]. Fragmentation was triggered for the top 15 peaks with charge 2–8 on the MS scan (data-dependent acquisition) with a 30 s dynamic exclusion window (normalized collision energy was 28). Precursors were isolated with a 1.4 m/z window and MS/MS spectra were acquired in profile mode with a resolution of 15,000 (maximum fill time of 80 ms, AGC target of 2e4 ions).

### Analysis of mass spectrometry data

Peptide and protein identification and quantification was done using the MaxQuant software (version 1.6.10.43) (Cox and Mann, 2008; Cox et al., 2011; Tyanova et al., 2016) and a *Saccharomyces cerevisiae* proteome database obtained from Uniprot. 10plex TMT was chosen in Reporter ion MS2 quantification, up to 2 tryptic miss-cleavages were allowed, protein N-terminal acetylation and Met oxidation were specified as variable modifications and Cys carbamidomethylation as fixed modification. The “Requantify” and “Second Peptides” options were deactivated. False discovery rate was set at 1% for peptides, proteins and sites, minimal peptide length was 7 amino acids.

The output files of MaxQuant were processed using the R programming language. Only proteins that were quantified with at least two unique peptides were considered for the analysis. Moreover, only proteins that were identified in at least two out of three MS runs were kept. A total of 3,550 proteins for the whole cell proteome and a total of 1,154 for the IP passed the quality control filters. Raw signal sums were cleaned for batch effects using limma (Ritchie et al., 2015) and further normalized using variance stabilization normalization (Huber et al., 2002). Proteins were tested for differential expression using the limma package for the indicated comparison of strains.

A reference list of yeast mitochondrial proteins was obtained from (Morgenstern et al., 2017). Gene set enrichment analysis was performed using Fisher’s exact test. A Benjamini-Hochberg procedure was used to account for multiple testing, where this was performed (Benjamini and Hochberg, 1995).

### Fluorescence microscopy

Manual microscopy was performed using Confocal Zeiss LSM 700 (Axio Examiner.D1) system or a Leica Dmi8 Thunder Imager. Images were acquired using a Plan-Apochromat 63x/1.4 Oil DIC M27 or an HC PL APO100x/1,44 Oil UV objective. Carl Zeiss with Immersol 518 F immersion oil (ne = 1.518) or Immersion Oil Type A 518 F, with wavelength of 488 nm (GFP) and the ZEN2009 or LAS X software. Further processing of images was performed in Fiji/ImageJ.

### Measurements of fluorescent intensity profiles

To visualize colocalization of GFP and RFP tagged proteins, fluorescent intensity profiles were measured. Therefore, images were acquired as described previously. Acquired images were imported into ZEN 3.3 (blue edition) software, and fluorescent intensity profiles were generated by drawing a line across the object to be analyzed (in this case across GFP or RFP foci).

Profiles could then be exported as TIF file.

### Colocalization analysis via Pearson’s Correlation Coefficient calculation

To provide a quantitative statement on the colocalization of RFP and GFP tagged proteins, the Pearson’s Correlation Coefficient was calculated. Therefore, images were acquired, and fluorescent intensity profiles were measured as described above. The obtained data table containing the distance in nm and the intensity values of each selected channel was exported as CSV file. Pearson’s Correlation Coefficient r was calculated as following:

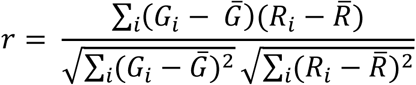

Gi and Ri refer to the intensity values of the green (488 nm) and red (561 nm) channels, respectively, of pixel i, whereas 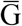 and 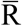 refer to the mean intensities of the red and green channels of each image. Pearson Correlation Coefficient can assume values between -1 and 1. A positive linear correlation results in a value of 1 and thereby describing perfectly linear related fluorescence intensities of two images. A negative linear correlation results in a value of -1, and no correlation results in a value of 0 (Dunn et al., 2011).

### Distance measurement of Hsp104-GFP foci

Characteristics of aggregates (Hsp104 foci) were quantified by measuring number and two-dimensional diameter. Therefore, microscopy images were acquired as described previously and imported into ZEN 3.3 (blue edition) software. Two-dimensional diameters were measured in nm by drawing a line across each Hsp104 foci. For quantification the number of aggregates per cell were plotted against the mean of two-dimensional aggregate diameter per cell.

### Cell cycle analysis by flow cytometry

For cell cycle analysis, cells were grown in selective lactate media. Clogger expression was induced for 4.5 h with 0.5% galactose. Cells were harvested (6,000 g, 1 min, RT) in the exponential phase. Cells were fixed by resuspending the pellet in 1 ml 70% EtOH and stored rolling over night at 4°C. Cells were washed with 1 ml H_2_O, the pellet stained with 500 µl FxCycleTM PI/RNase Staining Solution and stored in the dark for 30 min. Flow cytometry analysis was done using 2018 Attune NxT Flow Cytometer (Thermofisher Scientific) with an excitation of 535 nm. Data were then analysed with FlowJo 10.6.0 software (Tree Star).

## Notes

### Competing Interest Statement

The authors have declared no competing interest.

